# Defining characteristics of mesenchymal stem cell-derived matrix-bound nanovesicles compared to conditioned culture medium extracellular vesicles

**DOI:** 10.64898/2026.05.05.722048

**Authors:** Renata Dos Reis Marques, Maulee Sheth, Ava I. Salami, Supasek Kongsomros, Leyla Esfandiari, Marley J. Dewey

## Abstract

Matrix-bound nanovesicles (MBVs) are a type of small extracellular vesicle (EV) embedded in the extracellular matrix (ECM) throughout the body. MBVs have been previously isolated from various tissues and *in vitro-*cultured cell sheets, demonstrating remarkable attributes in regenerative medicine. However, differences between MBVs and conditioned culture medium-derived EVs (liquid-EVs) have yet to be characterized, and the field currently lacks specific protein markers that can identify MBVs from other EV subtypes. Here, we isolate MBVs and liquid-EVs from bone marrow mesenchymal stem cell (MSC) sheets and define differences in size, protein, and zeta potential between these EVs. We show that there is a correlation between cell-driven ECM deposition and MBV and liquid-EV production. We also find that MBVs are smaller, contain less protein per particle, and possess lower zeta potential than liquid-EVs. Interestingly, MBVs also comprise a distinct tetraspanin profile compared to liquid-EVs, with MBVs containing more CD63 and little to no CD81. Finally, we define that CD63, LAMP1, Alix, ITGβ1, and GRP94 and their abundance, may be markers specifically used to identify MBVs from liquid-EVs. Our study paves the way for the characteristic differentiation between MBVs from liquid-EVs, elucidates their differences in biogenesis, and reveals a potential connection between EV and ECM production.

## 1. Introduction

Extracellular vesicles (EVs) were first reported by Chargaff and West in 1946, described as “minute breakdown products” of blood cells recovered at a high centrifugal speed, but this study lacked any further investigation of their identity or role^1^. Decades later, EVs were formally defined, and several subpopulations of EVs have been identified across different species with different biological roles and characteristics, including microvesicles, exosomes, and apoptotic bodies, among others. EVs are known to play a pivotal role in several biological processes, such as wound repair^2–4^, cancer^5^, and aging^6^, serving as messengers between cell populations. Unfortunately, the classification of specific subtypes of EVs remains a challenge owing to their inherent structural and molecular heterogeneity. The International Society of Extracellular Vesicles (ISEV) has published adaptive recommendations for the categorization, isolation, and validation of EVs, yet these guidelines are context-dependent and contain limited information on solid tissue-derived EVs or matrix-associated EVs^7^.

Matrix-associated EVs are a subtype of EV residing in the extracellular matrix (ECM) and include migrasomes, matrix vesicles, and matrix-bound nanovesicles, among others^8^. Currently, little is understood about the production, specific markers, and role of these vesicles in the ECM, motivating their comprehensive characterization and categorization. Specifically, in this study, we aim to characterize matrix-bound nanovesicles (MBVs) and compare these to the most commonly studied class of EV: conditioned culture medium-derived EVs (liquid-EVs). Matrix-bound nanovesicles differ from other types of EVs in their native, matrix-bound state, as well as their harvesting methods. For example, liquid-EVs are released into biofluids (saliva, urine, blood, culture medium, etc.) and are widely thought to mediate cell-cell communication by encapsulation and exchange of signaling molecules^9^. In contrast, MBVs are isolated by digesting the extracellular matrix of whole tissues or *in vitro* cultured cell sheets^10^, and have pronounced immunomodulatory features^11^. This digestion process allows for MBVs entrapped within the ECM to be released, and these MBVs can then be retrieved using conventional EV isolation procedures^12^. MBVs can be isolated from non-mineralized tissues and ECM throughout the body, including the bladder, small intestine, heart, and muscle, among many others, and contain cargo distinctive to their original tissue location^13^.

Beyond isolation differences between MBVs and other types of EVs, MBVs also possess unique cargo and therapeutic properties. In a study comparing NIH 3T3 fibroblast conditioned medium-derived EVs to MBVs, MBVs lacked some common tetraspanins (CD9, CD63, CD81), but contained microRNAs unique to organ development and distinctly different lipid compositions^14^. MBVs have also been compared to liquid-EVs and calcifying matrix vesicles, a type of matrix-associated EV residing within mineralized tissues (calcified cartilage, bone). MBVs lacked characteristic mineral markers found in matrix vesicles (Annexin V, TNAP), and again, were missing some of the most common EV markers (CD63, Alix)^15^. The authors likewise found that MBVs had distinct lipid composition, microRNA cargo, and cytokine cargo compared to liquid-EVs and matrix vesicles. A more extensive proteomic characterization of mesenchymal stem cell (MSC) MBVs compared to their corresponding liquid-EVs demonstrated that MBVs were enriched in proteins related to enzymatic pathways, and MBVs had a greater angiogenic potential than liquid-EVs in an MSC and endothelial cell 3D co-culture model^16^. These works have established some characteristic and functional differences between MBVs and other vesicles, setting some of the foundations for identifying MBVs as a unique subpopulation of EV, yet there is a need for more comprehensive comparisons between MBVs and other EV subtypes.

The unique features of MBVs have led to major applications of MBVs towards regenerative medicine and wound repair. For example, MBVs have been used as a therapy for optic nerve repair ^17,18^, rheumatoid arthritis^19^, inflammatory bone diseases such as osteolysis^20^, among others. Outside of their role in tissue repair, MBVs also reflect the diseased state of tissues in their cargo, as MBVs isolated from diseased heart tissues contained several markers of heart disease and had less of a pro-healing impact on macrophages than their non-diseased counterparts^21^. Additionally, cancerous tissue contains MBVs that can be used to target cancer cells^22^, and aged tissues contain MBVs that can influence cancer progression^6,23^. These studies all indicate the potential to use MBVs as regenerative medicine therapeutics, as well as diagnostics and treatments for cancer and disease. As such, we find that there is a growing need for the specific delineation of EV identity and characteristics, given their recent rise in popularity in basic science research and translational applications.

Due to the beneficial properties of MBVs, a better understanding of their characteristics is critical for their effective use in clinical applications. Yet, while liquid-EVs and matrix vesicles are relatively well characterized in the field, the characteristics, biogenesis, and role of MBVs remain critically understudied. Specifically, the current MBV research landscape lacks information on MBV biomarker expression, such as tetraspanins, and more importantly, lack defining features of MBVs that distinguish these from other populations of EVs^14^. Thus, defining such features is crucial for improving MBV isolation protocols and understanding the purpose of MBVs in tissues.

Given unresolved ambiguities in MBV identification, we seek to compare the properties of liquid-EVs and MBVs to further elucidate differences in EV identity and report novel biomarkers that distinguish these two EV subpopulations on a molecular level. Here, we induce *in vitro* extracellular matrix deposition in three different patient populations of human bone marrow MSCs and isolate liquid-EVs and MBVs secreted from these cultures. We select MSCs as our cells of interest as EVs from these cells are promising cell-free therapies, and liquid-EVs from MSCs have already demonstrated prior therapeutic potential^2,3,24^. We also assess three patient donors (2 male, 1 female) of MSCs to determine unique characteristics of MBVs to better evaluate differences between liquid-EVs and MBVs. We assess key characteristics of EVs, including size, morphology, zeta potential, and protein content, comparing both donor-to-donor variability across the same EV subpopulation as well as differences between EV subpopulations consistent across all donors.

## 2. Materials and Methods

### 2.1. Cell culture

Three different donors of primary human bone marrow mesenchymal stem cells were purchased from RoosterBio® (Frederick, Maryland, USA) at passage 1 (**Table 1**). Cells were initially expanded in RoosterNourish™-MSC medium (RoosterBio Inc, K82003) until passage 3. On the fourth passage, cell culture medium was exchanged for a 1:1 mixture of RoosterNourish medium and low glucose DMEM (Gibco, 11885092) supplemented with 10% fetal bovine serum (Genesee, 25-514) and 1% antibiotic-antimycotic (Gibco, 15-240-062) to slowly acclimate cell cultures to normal growth medium. Cells were cultured in complete DMEM for all following passages (max. passage 6). Normal growth medium was exchanged every 3 days. All cultures were maintained at 37°C with 5% CO_2_.

**Table 1.** Summary of mesenchymal stem cell populations used in this study by age, sex, and corresponding lot number. Identifier column indicates naming convention of the three biological replicates used in this study.

| Cell type | Lot number | Sex | Age | Identifier |
| --- | --- | --- | --- | --- |
| Human bone marrow mesenchymal stem cells | 310305 | Female | 20 | 20F |
| Human bone marrow mesenchymal stem cells | 310307 | Male | 19 | 19M |
| Human bone marrow mesenchymal stem cells | 310310 | Male | 25 | 25M |

### 2.2. Cell sheet formation

Mesenchymal stem cells at passage 6 were seeded at an initial seeding density of 10,000 cells/cm^2^ in 150 mm petri dishes (Genesee Scientific, 25-203). Two days later, normal growth medium was supplemented with 50 μM L-ascorbic acid-2-phosphate sesquimagnesium salt hydrate (Tokyo Chemical Industry, A2521) to induce cell-driven extracellular matrix deposition *in vitro* and cells were cultured for an additional 10 days. On the ninth day of culture, growth medium was supplemented with extracellular vesicle-depleted fetal bovine serum to reduce cross-contamination of extracellular vesicles in the conditioned culture medium. Extracellular vesicle-depleted fetal bovine serum was prepared in advance by ultracentrifugation at 100,000*g* for 18 hours at 4°C. Cell sheet cultures were terminated on the twelfth day of culture for downstream experiments.

### 2.3. Scanning electron microscopy of cell sheets

Mesenchymal stem cell sheets were cultured in a coverslip-bottom petri dish (Mattek, P35G-1.5-14-C) as previously described. On the twelfth day of culture, cells were fixed with 10% formalin (Sigma, HT501128) for 10 minutes, washed three times in 1X PBS, and stored at 4°C until further experiments. Cell sheets were washed with ultrapure water then dehydrated in a series of ethanol solutions of increasing concentration (30%, 50%, 70%, 85%, 95%, and 100% ethanol for 5 minutes each). The cells were briefly washed twice in a 1:1 solution of hexamethyldisilazane (HMDS, Sigma, 379212) and 100% ethanol, then washed once in HMDS. Fresh HMDS solution was added to the cell sheets and allowed to evaporate overnight in a chemical hood. Dehydrated cell sheet samples were sputter coated with gold nanoparticles for 30 seconds using a SuPro Instruments ISC 150T Ion Sputter Coater. Samples were imaged using a Thermo Scientific Apreo C LoVac scanning electron microscope at 4kV and 13pA.

### 2.4. Sirius Red staining and quantitative assay

Mesenchymal stem cell sheets were cultured in 12-well plates as previously described in Section 2.2. On the twelfth day of culture, cells were fixed with 10% formalin for 10 minutes, washed three times in 1X PBS, and stored at 4°C until staining. Sirius red staining and subsequent stain quantification were performed as described by Szász et al^25^. A 0.1 w/v% solution of Direct Red 80 (Sigma, 365548) in 1% acetic acid was prepared in advance of staining. Fixed cell sheets were stained in Sirius Red solution for 1 hour at room temperature with shaking. Stained cell sheets were then rinsed in 0.1M HCl until no excess stain was observed. After this wash, cells were imaged using a Zeiss Axio Vert.A1 coupled to a CMOS microscope camera (AmScope, MU1803). The stain was eluted with 0.1M NaOH and absorbance was measured at 540 nm using a TECAN Infinite M Plex plate reader.

### 2.5. Cell sheet differentiation

Mesenchymal stem cells were seeded in each well of a 12-well plate at a seeding density of 10,000 cells/cm^2^ and cultured in L-ascorbic acid-2-phosphate as previously described in Section 2.2 to induce cell sheet formation. On the last day of culture, extracellular vesicle-depleted growth medium containing L-ascorbic acid was exchanged for osteogenic or adipogenic differentiation induction medium. Osteogenic differentiation was accomplished by supplementing normal growth medium with 50 μM L-ascorbic acid-2-phosphate, 10 mM β-glycerophosphate (Sigma, G9422), 100 nM dexamethasone (Sigma, D4902), and 2 mM Glutamax (Gibco, 35050061). Adipogenic differentiation of cell sheets was induced by exchanging growth medium with high glucose DMEM (Gibco, 11995-065) supplemented with 10% fetal bovine serum, 1% antibiotic-antimycotic, 10 μg/mL human recombinant insulin (Gibco, 12585014), 500 μM 3-isobutyl-1-methylxanthine (Fisher, AC228420010), 1 μM dexamethasone, and 200 μM indomethacin (Fisher, AC458030050). After three days of culture in adipogenic induction medium, growth medium was exchanged for adipogenic maintenance medium comprising high glucose DMEM supplemented with 10% fetal bovine serum, 1% antibiotic-antimycotic, and 10 μg/mL human recombinant insulin for two days. Control cell sheets were cultured in identical medium compositions without induction factors. All cultures were maintained for a total of two weeks, then fixed in 10% formalin for 10 minutes at room temperature for downstream assays.

### 2.6. Alizarin Red S staining and quantitative assay

Fixed osteogenic cell sheets were incubated for 10 minutes in a 0.5 w/v% solution of Alizarin Red S pH = 4.17 (Sigma, A5533). The Alizarin stain was vigorously washed with ultrapure water until no excess stain was observed. Stained cell sheets were stored at 4°C in ultrapure water until imaging. Stained osteogenic cell sheets were imaged using a Zeiss Axio Vert.A1 coupled to a CMOS microscope camera (AmScope, MU1803). For dye elution and quantification, a solution of 10 w/v% cetylperidinium chloride (Sigma, C0732) was added to stained cell sheets and incubated for 20 minutes. The absorbance of the eluted dye was measured at 570 nm using a TECAN Infinite M Plex plate reader.

### 2.7. Oil Red O staining and quantitative assay

Fixed adipogenic cell sheets were washed twice in ultrapure water and incubated in 60% isopropyl alcohol for 5 minutes. A stock solution of 3 mg/mL Oil Red O (Sigma, O0625) in 100% isopropyl alcohol was diluted to a working concentration of 1.8 mg/mL with ultrapure water, incubated for 10 minutes at room temperature, and filtered through a 0.22-µm PES membrane (CELLTREAT, 229747) immediately prior to staining. Cells were stained in working Oil Red O solution for 20 minutes at room temperature and washed vigorously with ultrapure water until no excess stain was observed. Stained adipogenic cell sheets were imaged using a Zeiss Axio Vert.A1 coupled to a CMOS microscope camera (AmScope, MU1803). Following this, stained cell sheets were incubated in isopropyl alcohol to elute Oil Red O stain and the absorbance of the eluted dye was measured at 492 nm using a TECAN Infinite M Plex plate reader for quantification.

### 2.8. Isolation of liquid-EVs and matrix-bound nanovesicles from mesenchymal stem cell sheets

Conditioned culture medium from the last three days of cell sheet cultures was collected to isolate liquid-EVs. Cell sheets were washed in 1X PBS twice to remove residual culture medium and the remaining cell-deposited extracellular matrix was digested with sterile-filtered 10 μg/mL Liberase TH enzyme (Sigma, 5401151001) in liberase buffer (50 mM Tris-HCl pH = 8.0, 200 mM NaCl, 5 mM CaCl_2_) for 3 hours at 37C with constant rotation to release matrix-bound extracellular vesicles. The collected conditioned culture medium and extracellular matrix digests were separately differentially centrifuged at 500*g* for 10 minutes, 2,500*g* for 20 minutes, and 10,000*g* for 30 minutes at 4°C. The remaining supernatants were filtered through a 0.22-μm PES membrane (Fisher, FB12566508) to remove large particles and concentrated using 100kDa molecular weight cut-off regenerated cellulose membrane ultra centrifugal filters (Sigma, UFC910024). Enriched extracellular vesicle samples were further purified using size-exclusion chromatography. Briefly, disposable chromatography columns (BioRad, 7321010) were loaded with 10 mL of Sepharose CL-2B beads (Sigma, CL2B300). Packed columns were flushed with 0.22-μm filtered 1X PBS and 1 mL of sample was loaded into the column at a time. Fractions 3 through 5 (out of 10 total fractions) were collected, pooled, and further concentrated using 100kDa filters. Purified extracellular vesicle samples were aliquoted and placed at -80°C for long-term storage. Extracellular vesicles were subject to no more than two freeze-thaw cycles. Columns were re-used for a total of five times with thorough cleaning between uses. For cleaning, columns were flushed with 1X PBS then cleaning solution (0.5M NaCl, 0.1M NaOH). Finally, columns were rinsed with ultrapure water until the pH of the flow-through was neutral, then equilibrated in 1X PBS. Columns were stored at 4°C in 1X PBS between uses.

### 2.9. Nanoparticle tracking analysis

Liquid-EV and matrix-bound nanovesicle samples were diluted 1:1,000 and 1:10,000 in ultrapure water, respectively. Diluted extracellular vesicles were analyzed using a Malvern Panalytical Nanosight NS300 at camera level 11, syringe pump speed 100, and detection level 5. All samples were analyzed immediately after isolation.

### 2.10. Transmission electron microscopy

Four microliters of extracellular vesicle samples were pipetted onto the carbon-coated side of a formvar/carbon film square 200-mesh copper grids (Electron Microscopy Sciences, FCF200CU50) and immediately wicked away using Whatman no. 1 filter paper (Cytiva, 1001-0155). This process was repeated once. Following this, grids were stained with four microliters of uranyl acetate and immediately blotted with filter paper twice. Mounted and stained grids were allowed to dry until imaging. Extracellular vesicle samples were imaged using a JEOL 1230 transmission electron microscope coupled with a Hamamatsu C4742-95 camera. Digital images were captured using AMT Image Capture (version 7.0.0.268).

### 2.11. Electrophoretic light scattering

Extracellular vesicle samples were diluted to 1 x 10^10^ p/mL in 1 mM PBS. Samples were loaded into a folded capillary zeta cell (Malvern, DTS1070) up to the top of the electrodes and inserted into the Malvern Zetasizer Nano ZS with the cell seam facing the front of the instrument. A total of five replicate measurements per sample were recorded with measurement sub-runs automatically determined by the Zetasizer software.

### 2.12. Isolation of whole cell lysate

Following cell sheet formation in a 6-well plate, cells were washed with 1X PBS and lysed in ice-cold RIPA buffer (Thermo Scientific, 89900) supplemented with phosphatase and protease inhibitors (Thermo Scientific, A32961) according to manufacturer instructions (1 tablet per 10 mL of lysis buffer). Whole cell lysate was scraped from culture dishes, collected, and incubated on ice with intermittent vortexing. Samples were aliquoted and stored at -80°C until protein quantification and downstream experiments.

### 2.13. Protein quantification

The protein concentration of whole cell lysates and extracellular vesicle samples was quantified using BCA (Thermo Scientific, 23227) and microBCA (Thermo Scientific, 23235) assays, respectively, according to manufacturer instructions. Whole cell lysates were diluted 1:10 in RIPA and extracellular vesicles were diluted 1:10 in 1X PBS prior to protein quantification. Bovine serum albumin standards were diluted in RIPA or 1X PBS to generate a standard curve. For both assays, multiple absorbance measurements (562 nm) of each well were performed and averaged using a TECAN Infinite M Plex plate reader.

### 2.14. SDS-PAGE

Using protein concentration values previously determined by BCA and microBCA assays, whole cell lysate and extracellular vesicle samples were combined with 4X Laemmli buffer (BioRad, 1610747) containing beta-mercaptoethanol (Sigma, M3148). Samples were then boiled at 95°C for 5 minutes and briefly centrifuged. Five micrograms of total protein were loaded into each lane of either a 10% or 12% Mini-PROTEAN TGX gel (BioRad, 4561034 and 4561043). The first lane of the gel was loaded with a pre-stained protein ladder as a molecular weight reference. Gel electrophoresis was performed in running buffer (25 mM Tris, 190 mM Glycine, 0.1% SDS) at 150V at room temperature. For Coomassie blue staining, a 12% SDS-PAGE gel was first incubated in 10% acetic acid, 50% ethanol, and 40% ultrapure water and microwaved for 30 seconds. After the gel was allowed to cool to room temperature, the solution was replaced with 0.001 w/v% Coomassie Blue R-250 in 7.5% acetic acid, 5% ethanol, and 87.5% water. The gel in staining solution was microwaved for 30 seconds then incubated for 1 hour with gentle rocking at room temperature. Finally, the gel was rinsed in ultrapure water to remove excess stain. The stained gel was imaged using a BioRad ChemiDoc MP.

### 2.15. Western blotting

Immediately after gel electrophoresis, separated proteins were transferred to a nitrocellulose membrane (BioRad, CAT# 1704158) using a BioRad Trans-Blot Turbo semi-dry transfer system. Pre-programmed transfer settings were selected based on target molecular weight (**Supplementary Table 1**). These membranes were then briefly washed in 1X PBS and incubated for 1 hour in blocking buffer (5 w/v% non-fat dry milk in 1X PBS) at room temperature with gentle shaking. Primary antibodies were diluted in 5 w/v% bovine serum albumin (Sigma, 10735078001) dissolved in 1X PBS-T (0.2%) according to **Supplementary Table 1**. Blots were incubated in primary antibody overnight at 4°C with gentle shaking. Following this, blots were washed vigorously for 5 minutes three times with 1X PBS-T (0.2%) to remove unbound antibody. In the meantime, Alexa Fluor 647 goat anti-rabbit secondary antibody (Invitrogen, A21244) was diluted 1:10,000 in 5 w/v% non-fat dry milk dissolved in 1X PBS-T. Blots were incubated in secondary antibody for 1 hour at room temperature protected from light with shaking. Lastly, blots were washed vigorously for 5 minutes three times with PBS-T. Prior to imaging, blots were rinsed in 1X PBS to remove residual Tween-20. Imaging was performed using a BioRad ChemiDoc MP.

### 2.16. Super-resolution imaging of EVs

Sub-diffraction limit resolution images of single EVs were obtained using a Nanoimager S Mark II microscope (Oxford Nanoimaging, Oxford, UK) equipped with a 100X, 1.4 NA oil immersion objective, an XYZ closed-loop piezo 736 stage, and dual or triple emission channels split at 640 and 555 nm. ∼1 x 10^10^ EVs in 1 mL were stained with a 5 μg/mL mix of antibodies, including anti-CD63-Cy38 (Oxford Nanoimaging, Oxford, UK), anti-CD81-AlexaFlour647 (Oxford Nanoimaging, Oxford, UK), and anti-CD9-Atto488 (Oxford Nanoimaging, Oxford, UK). Antibody mix-only samples were acquired as negative controls. Samples were processed according to the manufacturer’s instructions to immobilize the stained EVs on chips provided with the EV profiler kit. Eight fields of view were recorded for each sample using direct stochastical optical reconstruction microscopy (dSTORM). Analysis was performed using algorithms including filtering, drift correction, and DBScan clustering developed by ONI (Oxford Nanoimaging, Oxford, UK) via the Collaborative Discovery (CODI) platform.

### 2.17. Statistics

Statistics were performed using Prism software (GraphPad, Massachusetts, USA). Significant differences were indicated as p-values less than 0.05. For all data, assumptions of normality were checked using a Shapiro–Wilk test and equal variance with a Browne–Forsythe test. T-tests were performed for data with two groups, and ANOVA with a Tukey post-hoc for >2 groups. Other tests were instead used if assumptions of normality or equal variance were not met (i.e., Kruskal-Wallis for >2 group non-normal analysis). Error bars for all data are represented as mean ± standard deviation and n values are indicated in figure captions.

## 3. Results

### 3.1. MSCs deposit variable amounts of collagen in vitro and retain multipotency after cell sheet formation

After cell sheet formation, we first verified that MSCs secreted collagen into their ECM in the presence of L-ascorbic acid-2-phosphate. The matrices deposited by MSCs in culture (20-year-old female, 19-year-old male, and 25-year-old male) indeed comprised collagen as evidenced by staining with Sirius Red, a dye that ionically interacts with collagen molecules in acidic conditions (**Fig. 1A**). Intriguingly, collagen deposition significantly differed between MSC populations, with 25-year-old male (25M) cells depositing the most collagen and 19-year-old male (19M) cells the least (**Fig. 1B**). Fixed MSC sheets were additionally imaged using scanning electron microscopy to assess the presence of EVs in the ECM. Vesicle-like structures were observed in clusters on top of cells (**Fig. 1C**).

**Figure 1.**
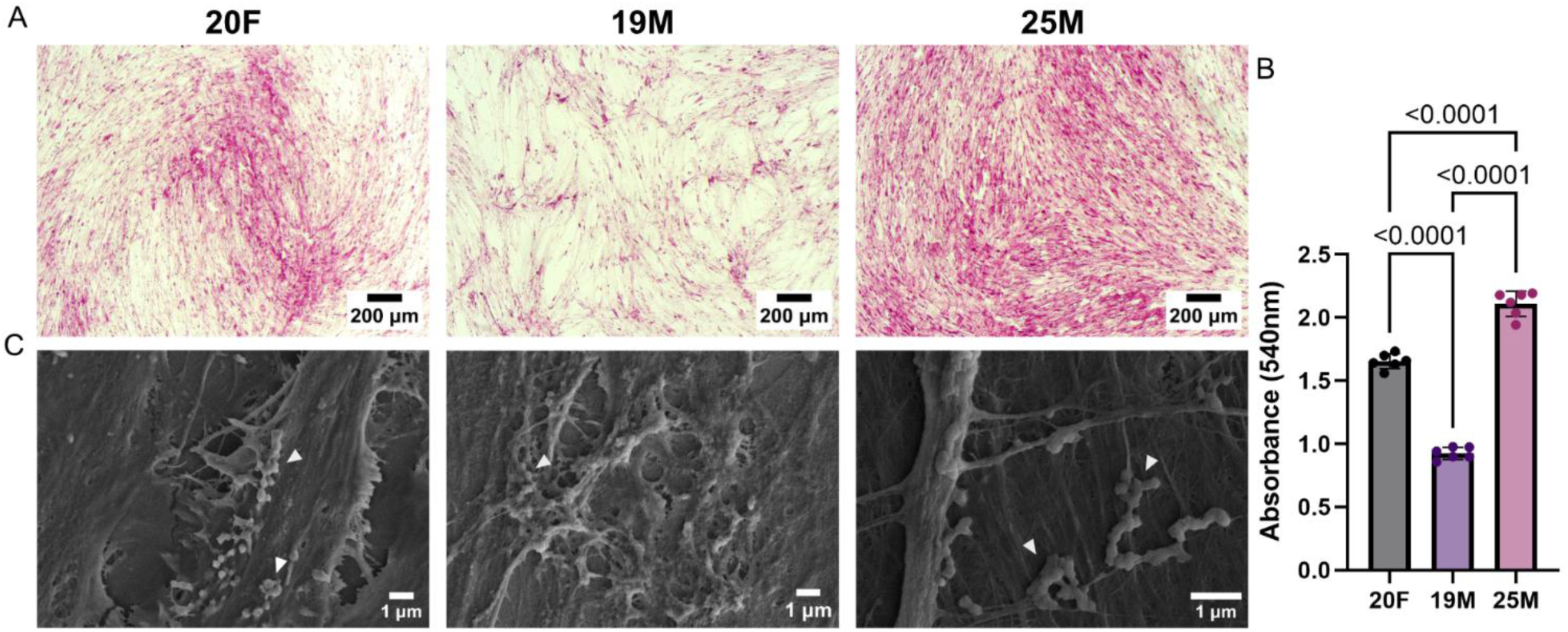
MSCs produce variable amounts of collagen *in vitro* in a donor-to-donor dependent manner and EV-like particles can be visualized within MSC sheets. (A) Representative images of Sirius Red-stained MSC sheets after 10 days of *in vitro* culture in L-ascorbic acid-2-phosphate. (B) Quantification of eluted Sirius Red dye from MSC sheets (n=6). Significance was calculated using a one-way ANOVA with a Tukey’s post-hoc test. (C) Representative SEM images of MSC sheets demonstrating vesicle-like structures on cells and ECM fibers. Potential MBV are indicated with white arrows.

On the last day of culture, we chemically induced osteogenic and adipogenic differentiation of MSCs and continued these cultures for an additional 14 days to allow for lineage commitment. We find that MSC sheets quantifiably maintain their multipotency after L-ascorbic acid-2-phosphate treatment and cell sheet formation as osteogenic and adipogenic cell sheets showed significant increases in mineralization and lipid droplet composition as measured by Alizarin Red S and Oil Red O staining, respectively, compared to non-treated groups (**Suppl. Fig. 1**). Notably, adipogenic 19M cell sheets contained less lipid droplets than other MSC populations (**Suppl. Fig. 1D**) and their osteogenic counterparts exhibited cell sheet peeling from the culture surface (**Suppl. Fig. 1B**).

### 3.2. MBVs are significantly smaller than liquid-EVs and MBV yield correlates to collagen deposition

Immediately following EV isolation, we performed nanoparticle tracking analysis to assess EV quantity and size. Across all donors, we found that liquid-EVs exhibited a wider size distribution than MBVs, though both EV subtypes were primarily within the 50-200 nm size range typically expected of small EVs (**Fig. 2**). On average, liquid-EVs possessed significantly larger mean particle diameter (162.1 ± 10.9 nm) than MBVs (117.8 ± 1.56 nm) (**Fig. 3A**), with liquid-EVs showing higher biological variability as evident by significant differences in particle size across different donors (**Fig. 3B**). In contrast, MBVs maintained similar mean particle size across different donors (**Fig. 3C**). Interestingly, 19M cells secreted higher quantities of liquid-EVs than other MSC populations, though this difference was only significant between 20F and 19M cells (**Fig. 3B**). On the other hand, 25M cells were the best producers of MBVs and 19M cells were the poorest (**Fig. 3C**) and this observed donor-to-donor trend in MBV yield was mirrored by that of collagen deposition (**Fig. 1B**). We were able to recover liquid-EVs and MBVs at an average of 3.65 x 10^10^ ± 9.58 x 10^9^ particles and 1.51 x 10^12^ ± 4.89 x 10^11^ particles, respectively, from three replicates per donor, representing a 41-fold difference in average MBV yield compared to liquid-EVs. Because our isolation times were different between groups, we cannot statistically compare yield differences between liquid-EVs and MBVs (**Fig 3B, C**). Both subclasses of EVs exhibit expected EV morphology with no notable differences upon inspection with transmission electron microscopy; however, there was a higher concentration of MBVs per field of view due to differences in particle concentration between EV preparations (**Fig. 2**).

**Figure 2.**
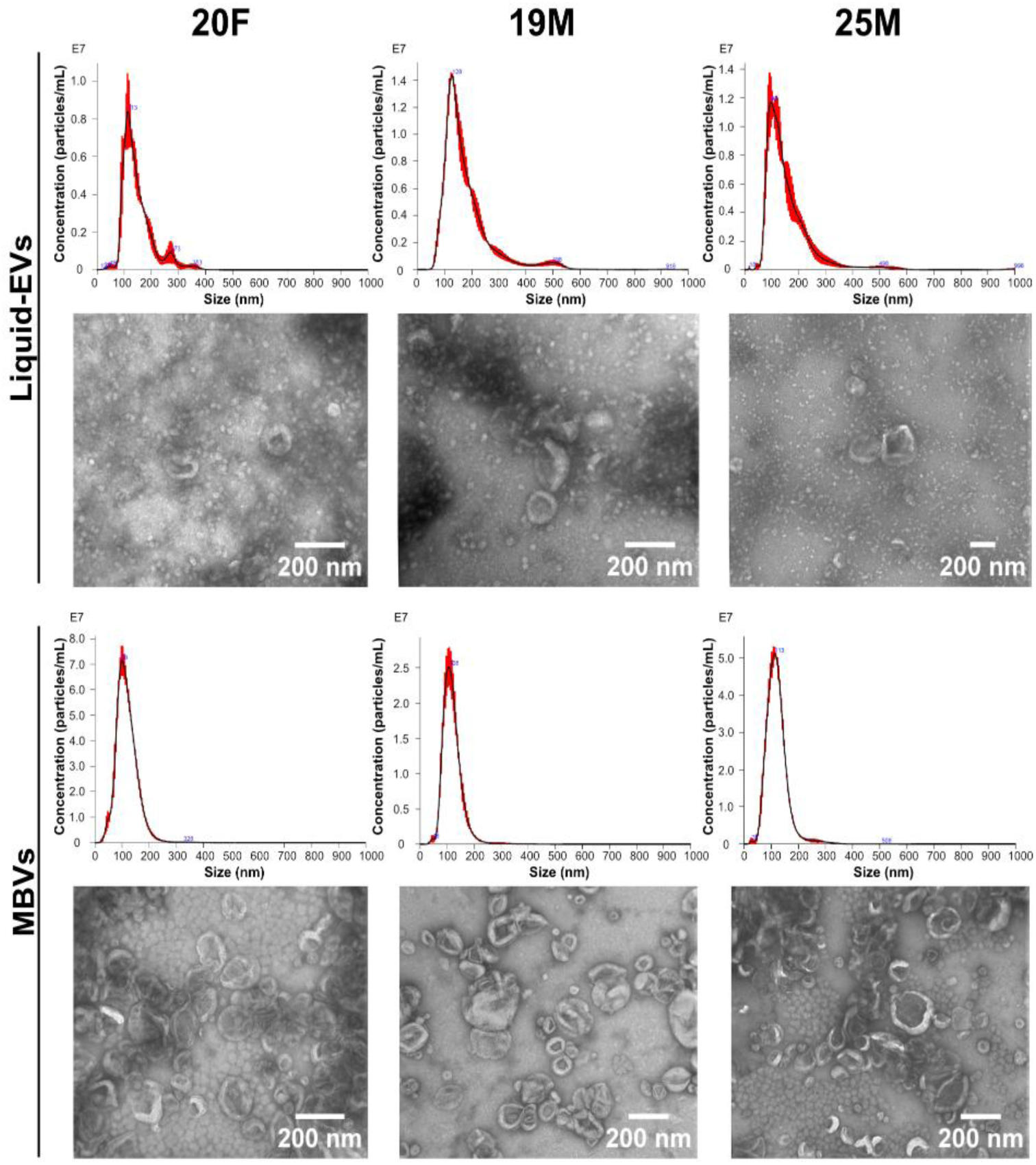
Liquid-EVs possess a wider size distribution than MBVs but both exhibit similar EV-like morphology. *Top:* Representative nanoparticle tracking analysis size distribution plots of liquid-EVs (1:1,000) and MBVs (1:10,000). Sample concentration in particles per milliliter (1 x 10^7^) plotted against size in nm. *Bottom:* Representative transmission electron microscopy images of liquid-EVs and MBVs.

**Figure 3.**
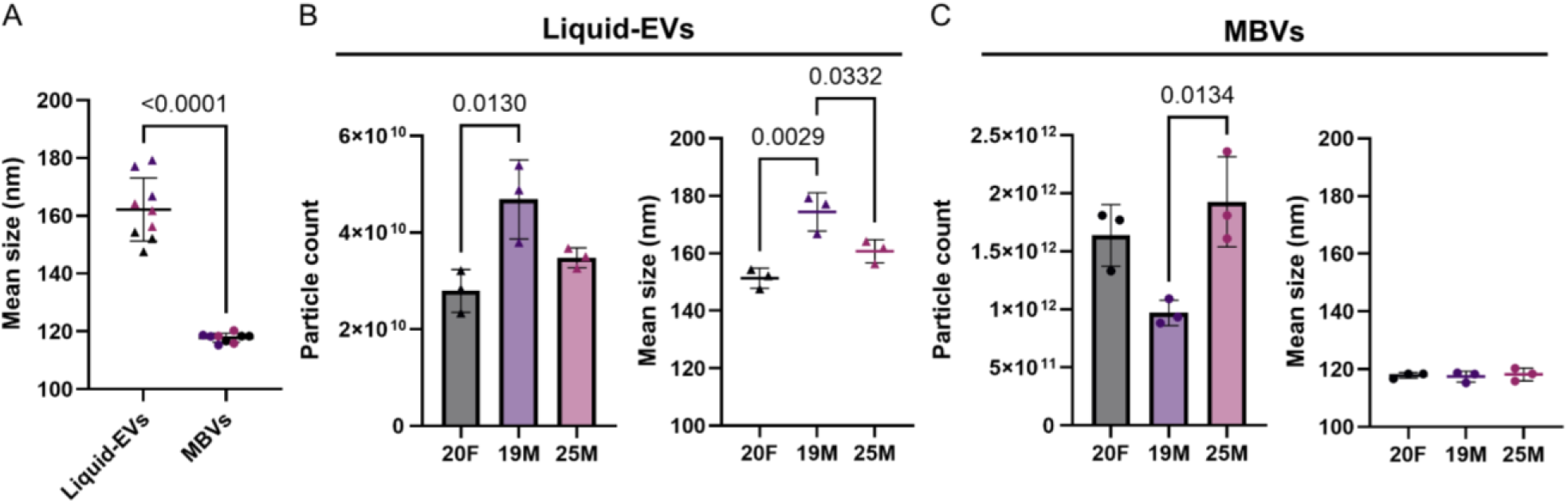
Liquid-EV and MBV yields differ between MSC donors and MBVs are smaller than liquid-EVs. (A) Comparison between liquid-EV and MBV mean size (n=3) (B) Quantification of liquid-EV yield and mean size (n=3). (C) Quantification of MBV yield and mean size (n=3). Significance was determined using a Welch’s t-test (A) and one-way ANOVA with Tukey’s post-hoc tests (B, C).

### 3.3. MBVs possess a more negative zeta potential and contain less protein per particle than liquid-EVs

The protein concentration of EVs was quantified using a microBCA assay and normalized to particle concentration for each sample to determine the amount of protein per particle. Intriguingly, liquid-EVs comprised significantly more proteins per particle than MBVs, with the former containing a combined average of 2.095 ± 0.2 femtogram of protein per particle compared to 0.042 ± 0.006 femtogram per particle measured in MBVs. Within each EV subtype, there were no notable differences in EV protein concentration across different donors (**Fig. 4A**). We additionally performed SDS-PAGE to assess differences in total protein content between EVs and whole cell lysates extracted from MSC sheets. While in this context SDS-PAGE does not provide information on specific proteins that may be contained within samples, we found that the distribution of total protein molecular weights is strikingly different between cells, liquid-EVs, and MBVs; however, these patterns were seemingly consistent between donors when compared within the same sample group (**Fig 4B**). Furthermore, electrophoretic light scattering revealed that the zeta potential of MBVs is significantly higher in magnitude (-42.5 ± 3.42 mV) than liquid-EVs (-33.3 ± 2.54 mV), though both of these EVs possessed net negative zeta potential values (**Fig. 4C**).

**Figure 4.**
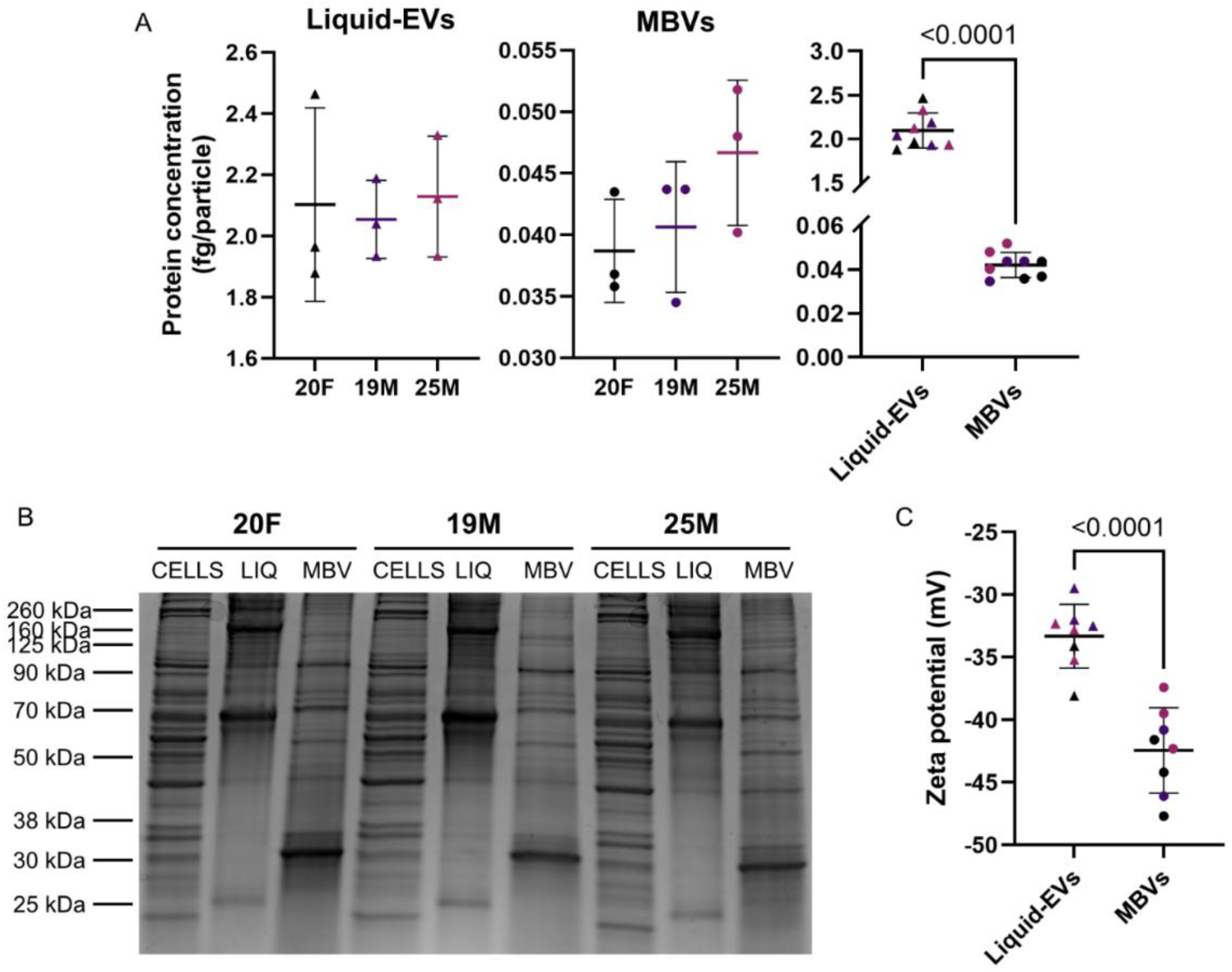
MBVs possess a more negative zeta potential and contain less protein than liquid-EVs. (A) Protein concentration of liquid-EVs, MBVs, and liquid-EVs compared to MBVs (N=3, n=3) in femtogram per particle. (B) SDS-page of whole cell lysates, liquid-EVs, and MBVs extracted from three different MSC donors. Band patterns differ between liquid-EVs, MBVs, and whole cell lysates, but are similar across biological replicates. (C) Zeta potential of liquid-EVs compared to MBVs from three different MSC donors (N=3). Significance was determined using a one-way ANOVA with Tukey’s post-hoc tests (A), Mann-Whitney test (A), and t-test (C).

### 3.4. MBVs are enriched in LAMP1, GRP94, CD63, and Integrin-β1 compared to liquid-EVs

A critical component of successful EV isolation validation includes the verification of molecular markers associated with EVs. ISEV’s most recent guidelines for minimal information in EV studies recommend the assessment of protein marker expression in EV samples that suggest the absence of non-EV co-isolates with an enrichment of EVs. This assessment often involves Western blot analysis of EVs and their respective whole cell lysates to probe protein expression across several different categories of proteins, including transmembrane proteins associated with the plasma membrane and endosomes, EV-associated cytosolic proteins, as well as components of non-EV co-isolates, to name a few^7^. As such, we sought to identify MBV-specific proteins that could differentiate MBVs from liquid-EVs and assess MBV sample purity as these markers have yet to be identified in the field. We performed Western blotting to assess markers related to ECM adhesion and EV biogenesis, as well as tetraspanins and putative negative EV markers. Of all protein markers tested, MBVs were consistently enriched for Integrin β1 (ITGβ1), lysosome-associated membrane protein 1 (LAMP1), ALG-2-interacting protein X (Alix), glucose-regulated protein 94 (GRP94), and CD63 when compared to liquid-EVs across all donors. In contrast, MBVs are devoid of CD81 unlike liquid-EVs. Expression of Integrin-αV (ITGαV) and heat shock protein family A member 13 (HSPA13) is low in both whole cell lysates and EVs (**Fig 5, Suppl. Fig. 3**).

**Figure 5.**
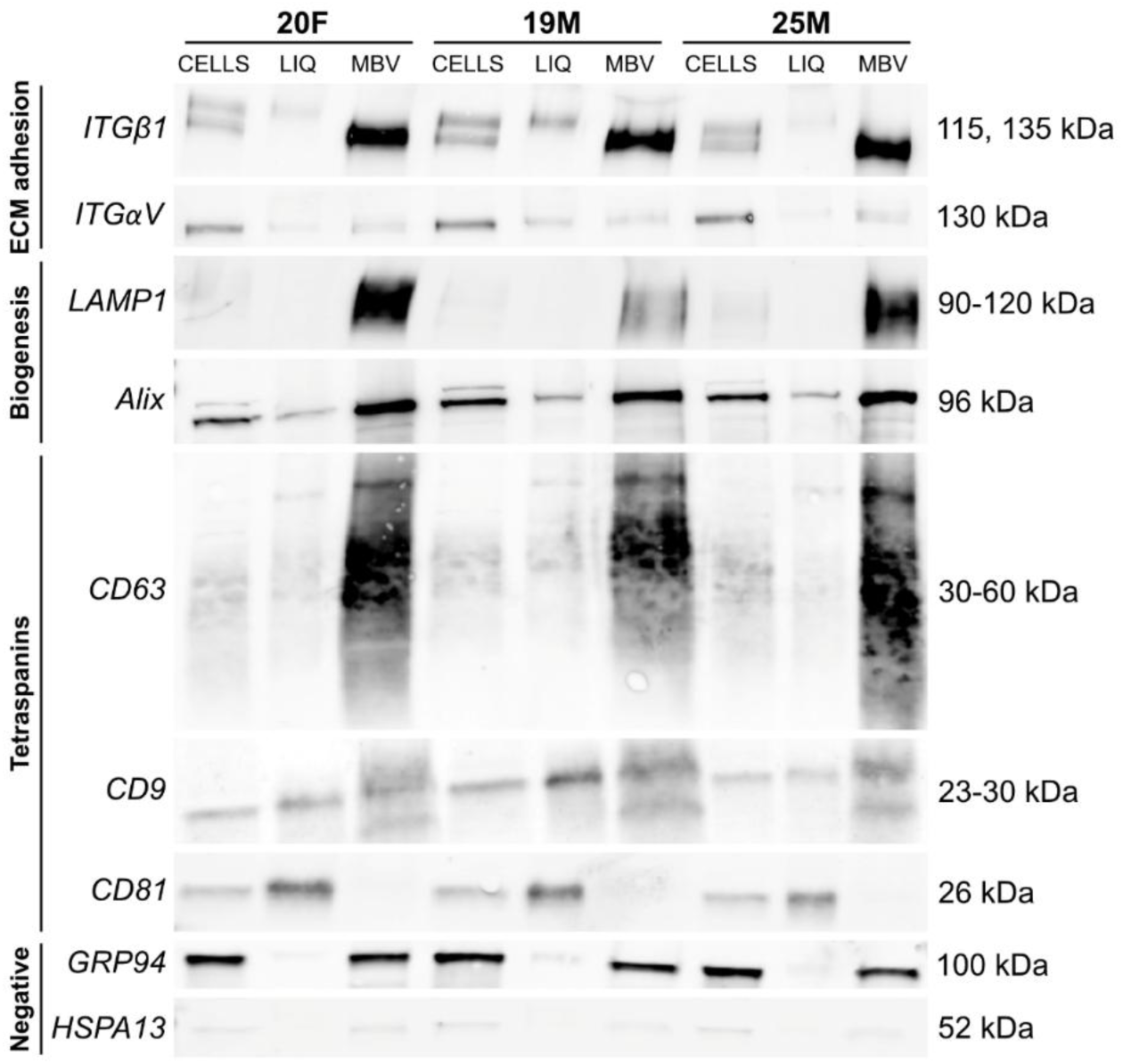
MBVs are enriched in ITGβ1, CD63, LAMP1, Alix, and GRP94 compared to liquid-EVs. Western blotting of MBVs compared to liquid-EVs (LIQ) and whole cell lysates (CELLS) from different MSC donors.

### 3.5. Super-resolution microscopy confirms enrichment of CD63 and lack of CD81 in MBVs relative to liquid-EVs

To confirm Western blotting results, we performed dSTORM super-resolution microscopy on liquid-EVs and MBVs isolated from MSCs. We assessed expression of three common tetraspanins, namely CD9, CD81, and CD63, at a single-particle level. Similar to our Western blot results, we found that MBVs were enriched for CD63 relative to liquid-EVs and lacked CD81 (**Fig. 6**). These differences were evident at a single-EV scale (**Fig. 6A**) as well as bulk quantification of over 1500 EVs per sample (**Fig. 6B, Supplemental Figure 2**). In particular, liquid-EVs exhibited a broader tetraspanin distribution across all three markers. A majority of the population were at least double-positive, and between 10-15% were triple-positive for all three markers. In contrast, MBVs demonstrated a more restricted marker profile, with a substantial 60-75% of vesicles single-positive for CD63 and 13-27% double-positive for CD63 and CD9, with minimal to no detectable CD81 expression. This confirmed that MBVs and liquid-EVs possessed distinct tetraspanin profiles.

**Figure 6.**
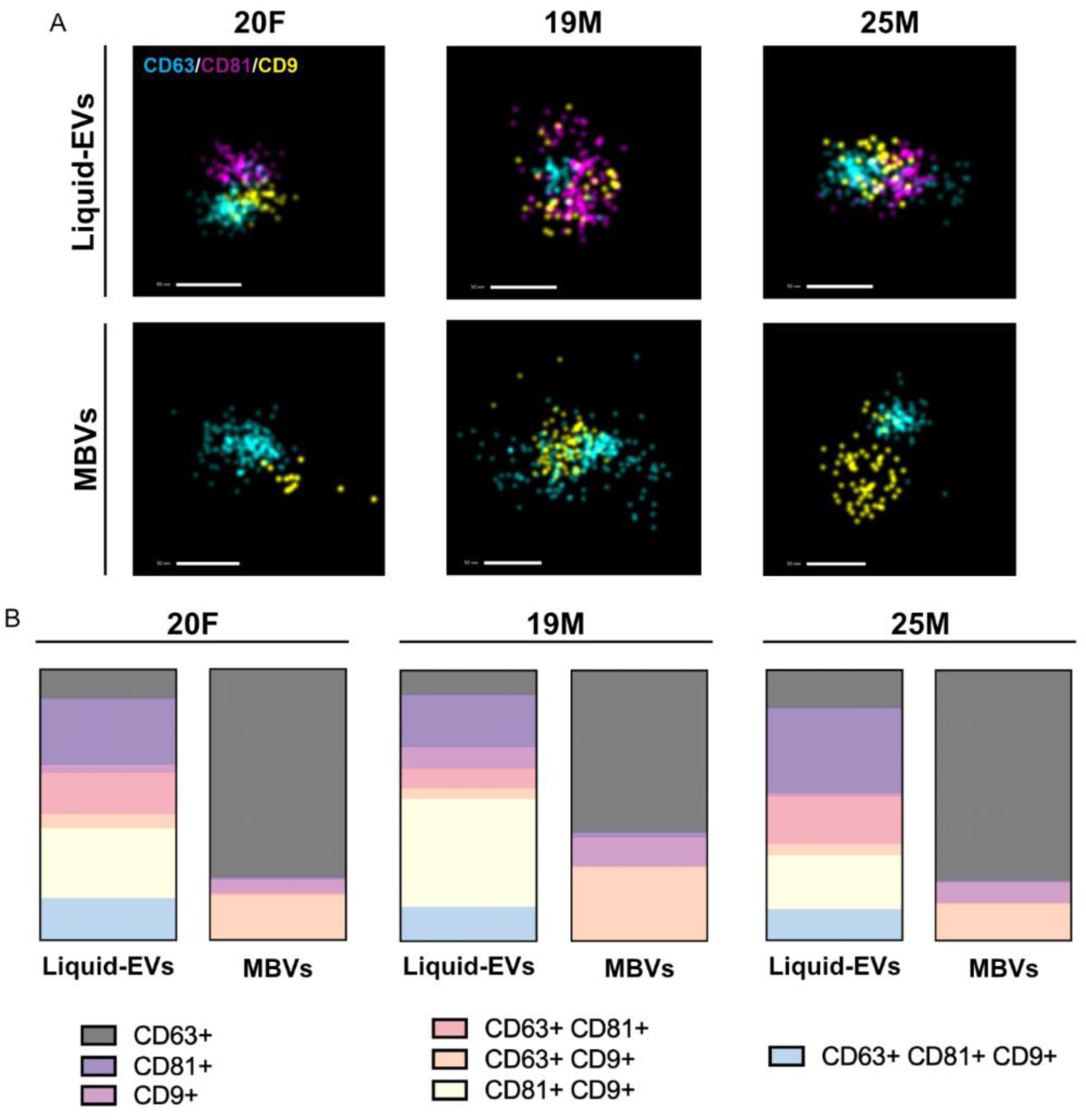
MBVs and liquid-EVs possess distinct tetraspanin expression profiles. (A) Representative dSTORM images of triple-positive liquid-EVs (top) and double-positive MBVs (bottom) from each donor. Samples were labeled with CD63 (Cy38, cyan), CD81 (AF647, magenta), and CD9 (Atto488, yellow). Scale bar = 50 nm. (B) Quantitative donor-specific tetraspanin distribution analysis between liquid-EV and MBV populations. A minimum of 1600 vesicles (MBVs or liquid-EVs) from at least three separate fields of view were analyzed per sample.

## 4. Discussion

MBVs have proven effective as therapeutics in a range of diseases and inflammatory tissue pathologies; however, the current MBV research landscape is limited by a poor understanding of MBV identifying characteristics that distinguish these from other types of EVs, such as those secreted into biofluids and conditioned culture medium (liquid-EVs). To address this critical gap in the field, we compared various EV properties, including size, zeta potential, and protein composition between MBVs and liquid-EVs extracted from the same cell populations and further defined such differences between these EV subpopulations. An additional strength of our study is the inclusion of biological heterogeneity by isolating EVs from three different human mesenchymal stem cell populations.

We first discovered significant (p < 0.05) differences in EV size and yield. Mean particle sizes were smaller in MBVs than liquid-EVs, which aligns with other published literature^26^ (**Fig. 3A**), and liquid-EVs uniquely demonstrated significant donor-to-donor differences in mean size (**Fig. 3B**). Intriguingly, the observed donor-to-donor trend in MBV yield was similar to that of collagen deposition, as 25M produced the greatest amount of collagen and the highest number of MBVs, and 19M the least amount of collagen and MBVs (**Fig. 3C and Fig. 1B**), suggesting a potential connection between ECM production and MBV secretion. Our results also indicated a potential trade-off between the extent of liquid-EV and MBV secretion as 19M cells produced the greatest number of liquid-EVs but the least MBVs, and 25M the greatest MBVs and least liquid-EVs (**Fig. 3B, C**). We also found a significantly higher yield of MBVs compared to liquid-EVs. This difference could be due to EV isolation times as liquid-EVs were isolated from the last 3 days of culture, whereas MBVs were isolated from the ECM that accumulated over the entire culture period. We decided to collect conditioned culture medium in the last 3 days of culture as the alternative would introduce a disparity in isolation times and/or freeze-thaw cycles between liquid-EVs and MBVs that may differentially alter EV heterogeneity and integrity. However, this finding is contrary to some published literature, which demonstrated a significantly reduced yield of MBVs compared to liquid-EVs from MSCs^26^. This could be due to a difference in isolation time-scales and methodologies, as this lower yield in MBVs is likely attributed to a much shorter culture of only 48 hours^26^ compared to our 12-day period, in which the former would result in less ECM deposition and less MBV production. Our results demonstrate that the entrapment of MBVs in ECM may be an advantageous strategy to increase EV yield compared to liquid-EVs, and some of the first results linking ECM production with MBV production.

Beyond EV quantity and size, we also found MBVs and liquid-EVs to be distinguished by their protein concentration and expression. We observed significant differences in protein concentration between liquid-EVs and MBVs across all three donors. Additional separation of total proteins by molecular weight (SDS-PAGE) revealed strikingly different distributions between cells, liquid-EVs, and MBVs; others have compared electrophoretically separated proteins from liquid-EVs and MBVs derived from 3T3 fibroblasts cultured *in vitro* and arrived at a similar conclusion^14^. To further narrow differences in protein composition between MBVs and liquid-EVs, we then assessed their expression of specific proteins associated with ECM adhesion, EV biogenesis, and non-EV co-isolated structures, some of which are listed in MISEV 2023^7^.

We examined the presence of GRP94 in our EV preparations as GRFP94 is most commonly used as a negative marker to indicate the absence of contaminating cellular debris in EV preparations. However, GRP94 has also been identified in microvesicles^27,28^ and larger vesicles^29^, as well as conditions of cellular stress^30^. In bone marrow MSCs, EVs have been shown to be devoid of GRP94^31^, though others have also shown that MSC EV expression of GRP94 is possible under severe hypoxia^32^. Indeed, our liquid-EVs lacked GRP94 as expected, though it remains uncertain why our MBVs expressed GRP94; this may be attributed to our lack of decellularization prior to MBV isolation, in which the ECM digestion process may cause cellular stress and apoptosis. However, we avoided the use of decellularization agents such as trypsin can also cause a mass release of EVs, which would introduce another challenge to the isolation and comparison of liquid-EVs to MBVs^33^.

Many of the other proteins we examined to be differentially expressed between liquid-EVs and MBVs are associated with EV biogenesis, such as LAMP1, Alix, and CD63. LAMP1 is associated with the lysosome and late endosome, as well as endo-lysosomal vesicles^34^. In HeLa cells, researchers reported that EVs enriched in both CD63 and LAMP1 could be used to identify exosomes from plasma membrane-derived EVs (i.e. ectosomes) that are also present in liquids^35^. Others have shown that matrix vesicles and adipose EVs can also express LAMP1 under certain conditions^36–38^. Our results strongly suggest that the biogenesis of MBVs is related to the endo-lysosomal system given a co-concomitant enrichment in LAMP1 and CD63; however, conclusions regarding the ubiquity of LAMP1 in MBVs, as well as claims regarding the endo-lysosomal biogenesis of MBVs, would require LAMP1 expression to be shown across MBVs from multiple cell types and direct evidence of endo-lysosomal involvement in MBV biogenesis. Beyond the role of LAMP1 in EV biogenesis, a recent study demonstrated that artificial vesicles engineered to display LAMP1 had “robust uptake” by HEK293FT cells^39^, further suggesting that the native presence of LAMP1 on MBVs may increase their uptake by cells relative to liquid-EVs. While various types of EVs have been previously shown to express LAMP1, we are the first to show that bone marrow MSC MBVs uniquely express LAMP1 compared to liquid-EVs, suggesting LAMP1 as a marker to distinguish between these two EV populations in MSCs. Additionally, to our knowledge, LAMP1 expression of any MSC EV subtype remains to be reported altogether. MBVs also exhibited higher expression of Alix compared to liquid-EVs, though both expressed Alix to some extent. In addition to serving as a widely reported EV protein marker, Alix is an accessory protein to endosomal sorting complexes required for transport (ESCRT) and has been shown to play roles in exosome biogenesis^40^, tetraspanin sorting and delivery^41^, and microRNA packaging into EVs^42,43^, among many others. In MSCs, the absence of Alix expression has a severe effect on small EV production; namely, a decrease in EV yield, supported by particle numbers, total protein concentration, and tetraspanin (CD81, CD9, CD63) expression, as well as a decrease in microRNAs per EV^43^. The differential expression of Alix in MSC liquid-EVs and MBVs in our studies further supports differences in biogenesis pathways between these two EV subclasses, including potential differences in microRNA recruitment into these EVs.

Much like GRP94 and Alix, tetraspanins represent some of the most popular protein markers used to identify EVs and distinguish EV subpopulations. Functionally, tetraspanins are also involved in EV biogenesis, cargo selection, and targeting and uptake^44^. For the first time, using super-resolution microscopy of EVs at a single-particle level and Western blotting of bulk samples, we showed that MBVs were primarily characterized by EV subpopulations expressing CD63, CD9, or both, whereas liquid-EVs comprised various combinations of CD63, CD9, and/or CD81. While the diverse MSC liquid-EV tetraspanin profiles including all three tetraspanins reported herein are supported by previous studies^45^, an enrichment of CD63 in MBVs contradicts some previous findings comparing 3T3 fibroblast liquid-EVs and MBVs, in which MBVs had a lower enrichment in CD63 than liquid-EVs^14^. This discrepancy in MBV CD63 expression could be due to differences in species (murine vs. human), cell type (fibroblasts vs. MSCs), and MBV isolation methods. Nonetheless, our results highlight the relevance of tetraspanin-agnostic EV isolation methods and the need to characterize EVs holistically, beyond tetraspanin presence, given the stark differences in tetraspanin enrichment between liquid-EVs and MBVs, and tetraspanin profiling could be used to identify MBVs by their lack of CD81.

While defining the biogenesis of MBVs is critical to understanding the production of these by cells, we also sought to elucidate some potential mechanisms behind the relationship between MBVs and the ECM. To begin to understand this relationship, we assessed the expression of Integrins β1 and αV, which could play a role in tethering MBVs to the matrix. EV-bound integrins are involved in cellular uptake^46–48^, and, most importantly, EV adhesion to ECM proteins. For instance, EVs isolated from the breast cancer cells express Integrin β1 and interact with nonfibrillar collagen via RGD-binding integrins^49^. Others have also shown specific, integrin-mediated interactions between EVs and ECM proteins, such that microvesicles bind to collagen I but not collagen III and fibrin, and collagen I binding is disrupted by EV treatment with RGD or an Integrin-α2β1 inhibitor^50^. We found that MSC MBVs are enriched in Integrin-β1 compared to their liquid-EV counterparts, and given these studies, it is plausible that MBV enrichment in Integrin-β1 relative to liquid-EVs explains the innate ability of the former to remain bound to ECM in tissues. Additionally, this could explain previous studies demonstrating a much longer *in vivo* retention of MBVs at the site of subcutaneous injection (7 days)^20,51^ compared to liquid-EVs, which often circulate out of the body within 24-48 hours^52^. Our results highlight Integrin β1 as a distinguishing protein that could be used to identify MBVs from other EVs and suggest that MBVs may bind better to the ECM due to an enrichment in Integrin β1 compared to liquid-EVs.

Finally, we assessed the zeta potential of MBVs compared to liquid-EVs to evaluate differences in colloidal stability, and such, the prevention of aggregation that may affect their efficacy. We found the zeta potential of MBVs to be significantly higher in magnitude than that of liquid-EVs. Particles with an absolute zeta potential greater than 30 mV are generally considered to be stable, with higher values indicating strong electrostatic repulsion between particles, further preventing aggregation. While both liquid-EVs and MBVs averaged below -30 mV, the significantly lower zeta potential of MBVs may indicate higher colloidal stability and a decrease in tendency for aggregation compared to liquid-EVs. This negative charge of MSC liquid-EVs is consistent with literature findings, although there is a difference in magnitude between our liquid-EVs and published literature (-33.3 mV vs. -17.9 mV, respectively)^53^; however, we are the first to report the zeta potential of MBVs and its distinction from the more well-characterized zeta potential of liquid-EVs.

## 5. Conclusion

In summary, we provide some of the first unique biomarkers and characteristics to identify mesenchymal stem cell MBVs from liquid-EVs. Notably, we discovered GRP94, LAMP1, Alix, CD63, and ITGβ1 are specifically enriched in MBVs and not liquid-EVs. The presence of LAMP1 and CD63 indicates MBV biogenesis may be of endo-lysosomal origin, and LAMP1 may promote better cellular uptake of MBVs than liquid-EVs based on other published findings. Furthermore, the enrichment of ITGβ1 may explain how MBVs bind to the ECM, and significantly, we find that as collagen production increases, quantities of MBVs increase while liquid-EVs decrease. This information lays the groundwork for the discovery of the biogenesis of MBVs, their role in ECM production, and verification of their isolation from other EV populations.

## Acknowledgements

This work is supported by funding from the National Institute of Health NIGMS (R35GM160216) (MJD) and the National Science Foundation NSF CAREER ECCS (2046037)(LE) and University of Cincinnati College of Engineering startup funds (LE). This work was also partially supported by a National Institutes of Health T32 Training Grant (1T32GM141846) (RDM). The content is solely the responsibility of the authors and does not necessarily represent the official views of the National Institutes of Health or National Science Foundation. This study utilized instrumentation provided by the University of California-Santa Barbara’s Biological Nanostructures Laboratory, the Neuroscience Research Institute-Department of Molecular, Cellular and Developmental Biology Microscopy Facility, and the Materials Microscopy and Microanalysis Facility. The authors would like to thank Drs. Carolyn Mills for granting our use of a BioRad ChemiDoc MP for the completion of this study.

## Author Contributions

**Renata Dos Reis Marques:** Methodology, Investigation, Formal analysis, Writing – Original Draft, Visualization. **Maulee Sheth:** Methodology, Investigation, Formal analysis, Writing – Original Draft, Visualization. **Ava Salami:** Investigation. **Supasek Kongsomros:** Methodology, Investigation, Formal analysis. **Leyla Esfandiari:** Writing – Review & Editing, Supervision, Project Administration, Funding acquisition. **Marley Dewey:** Conceptualization, Methodology, Writing – Review & Editing, Supervision, Project administration, Funding acquisition.

## Data Sharing

Data will be made available upon reasonable request.

