## Supplemental Information for "Defining characteristics of mesenchymal stem cell-derived matrix-bound nanovesicles compared to conditioned culture medium extracellular vesicles"

| **Target/Antibody** | **Vendor** | **Catalog number** | **Transfer setting** | **Antibody dilution** |
|---|---|---|---|---|
| LAMP1<br>Rabbit PolyAb | Proteintech | 21997-1-AP | 1 MINI TGX | 1:5,000 |
| Integrin Alpha V<br>Rabbit PolyAb | Proteintech | 27096-1-AP | 1 MINI TGX | 1:2,000 |
| Integrin Beta 1 Rabbit mAb | Cell Signaling Technology | 34971 | 1 MINI TGX | 1:1,000 |
| Alix Rabbit PolyAb | Proteintech | 12422-1-AP | 1 MINI TGX | 1:1,000 |
| CD63 Rabbit PolyAb | Proteintech | 25682-1-AP | 1 MINI TGX | 1:1,000 |
| GRP94 Rabbit PolyAb | Proteintech | 14700-1-AP | 1 MINI TGX | 1:5,000 |
| CD81 Rabbit mAb | AlgentBio | ABXB62284 | LOW MW | 1:1,000 |
| CD9 Rabbit PolyAb | Proteintech | 20597-1-AP | LOW MW | 1:1,000 |
| HSPA13 Rabbit PolyAb | Proteintech | 12667-2-AP | 1 MINI TGX | 1:1,000 |

**Supplementary Table 1.** Summary of protein targets, primary antibodies (type, vendor, catalog number, dilution factor), and transfer protocols.

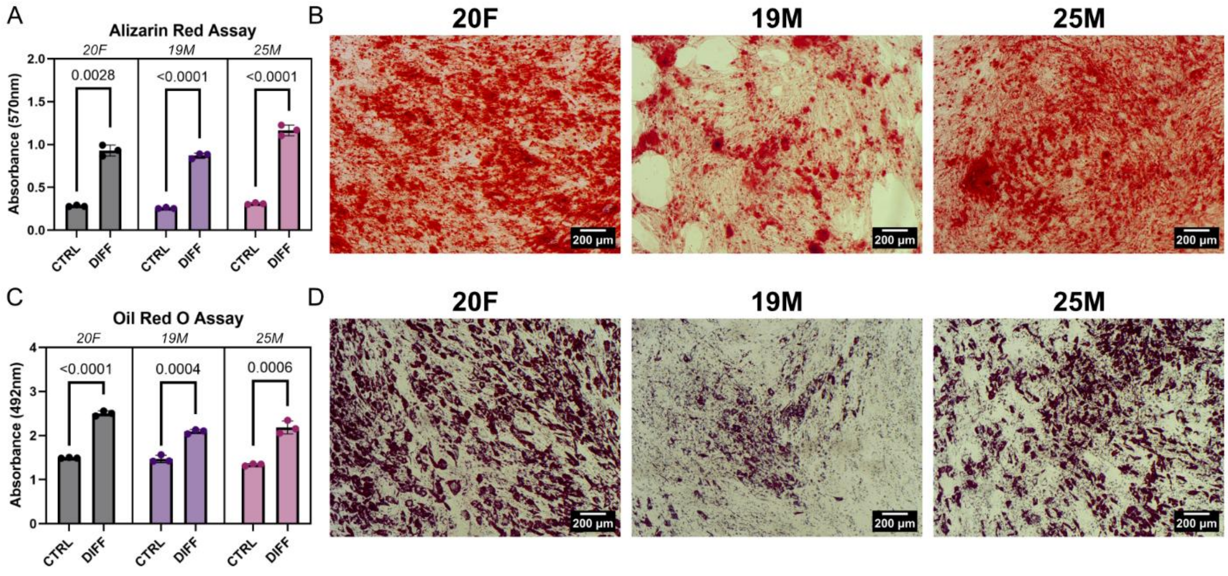


**Supplementary Figure 1. MSCs retain multipotency after cell sheet formation.** (A) Alizarin Red stain quantification of MSC sheets treated with and without osteogenic induction medium. (B) Representative images of Alizarin Red stained MSC sheets after osteogenic differentiation. (C) Oil Red O stain quantification of MSC sheets treated with and without adipogenic induction medium. (D) Representative images of Oil Red O stained MSC sheets after adipogenic differentiation. *CTRL* and *DIFF* denote undifferentiated and differentiated cultures, respectively.

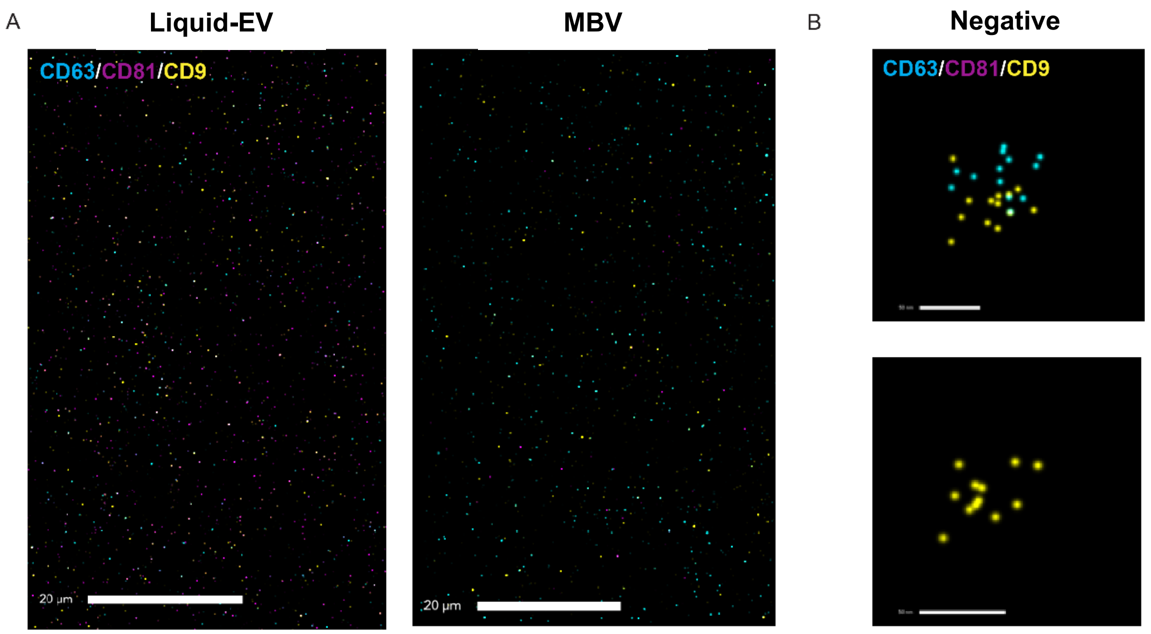


**Supplementary Figure 2.** (A) Representative dSTORM field of views showing captured MBVs or liquid-EVs. Scale bar = 20 µm. (B) Representative positive events for antibody mix-only negative control. Scale bar = 50 nm.

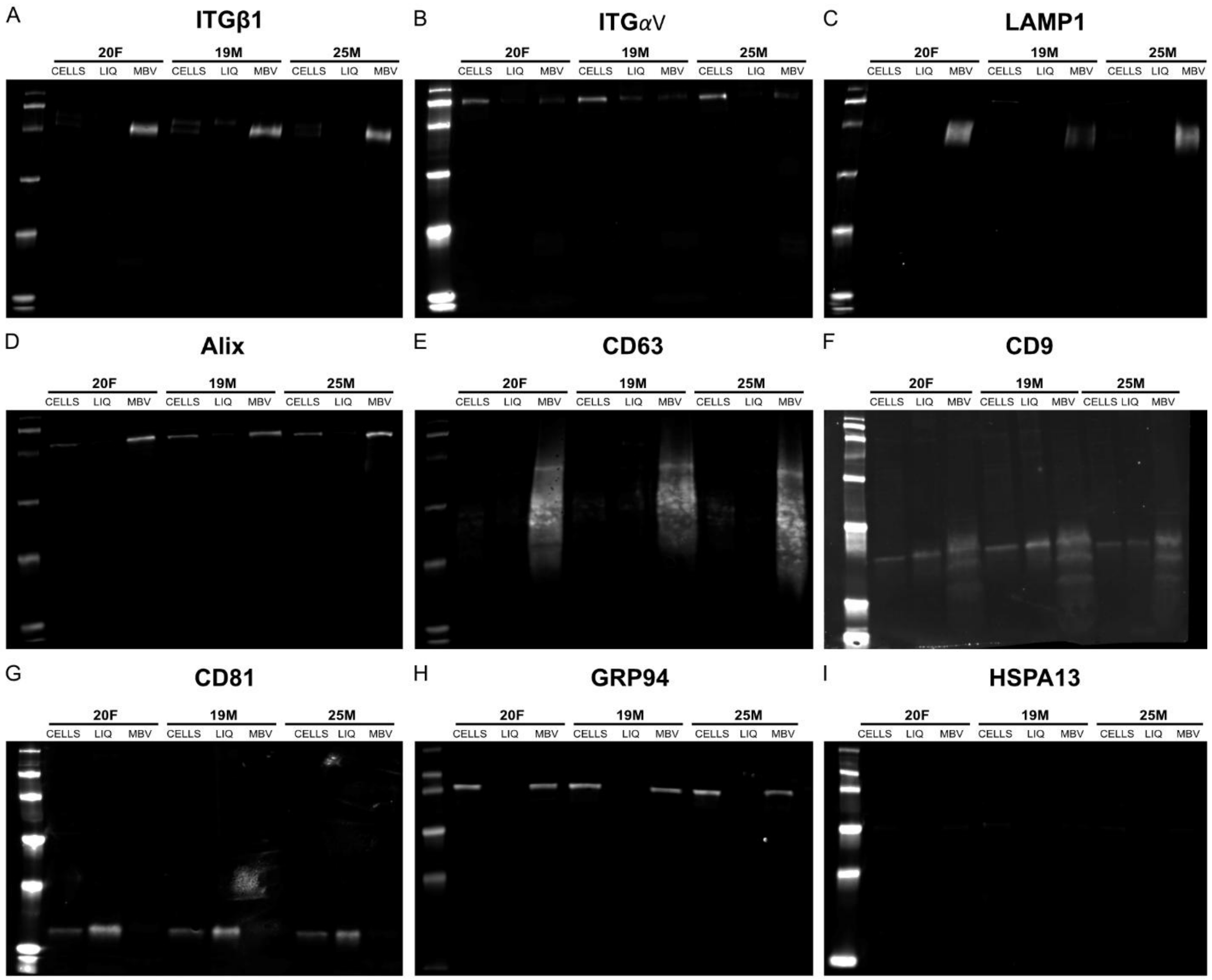


**Supplementary Figure 3.** Raw images of Western blots detecting MSC (CELL), liquid-EV (LIQ), and MBV expression of (A) Integrin β1, (B) Integrin αV, (C) Lysosome-associated membrane protein 1 (LAMP1), (D) ALG-2-interacting protein X (Alix), (E) CD63, (F) CD9, (G) CD81, (H) Glucose regulated protein 94 (GRP94), and (I) Heat shock protein family A member 13 (HSPA13).
